# Characterizing the Conformational Dynamics of an Intrinsically Disordered Localization Sequence

**DOI:** 10.64898/2026.03.04.709735

**Authors:** Matthew Brownd, Parth Chaturvedi, Ashkan Fakharzadeh, Mahmoud Moradi

## Abstract

Mitochondrial localization peptides (MLPs) play a critical role in directing proteins to mitochondria, yet how subtle sequence variations influence their conformational behavior remains poorly understood. Here, we investigate the conformational dynamics of a 15-residue MLP derived from the androgen receptor, together with a comprehensive panel of single-residue variants generated by systematic substitution at the second position. Across all variants, the peptide remains intrinsically disordered, exhibiting broad conformational heterogeneity and no stable folded state. Global measures of compactness show that single-residue substitutions induce only modest changes to overall peptide dimensions. In contrast, residue-level analysis reveals that the identity of the second residue subtly reshapes local structural preferences, particularly near the N-terminus. Small or hydrophobic substitutions enhance transient *α*-helical sampling, whereas polar or charged substitutions favor increased disorder and *β*- or polyproline-like conformations. Comparison across variants further distinguishes mutations that preserve wild-type-like structural behavior from those that produce more pronounced deviations in the conformational ensemble. Enhanced sampling simulations highlight the complexity and ruggedness of the underlying free-energy landscape and demonstrate the challenges associated with achieving convergence for short intrinsically disordered peptides. Collectively, these results show that even minimal sequence changes can bias the dynamic structural ensemble of mitochondrial localization peptides, suggesting a potential mechanism by which targeting efficiency may be modulated. More broadly, this work underscores the importance of advanced sampling strategies for accurately characterizing intrinsically disordered localization signals and provides a framework for connecting sequence variation to functional targeting behavior.

## Introduction

The androgen receptor (AR) is a member of the nuclear receptor superfamily, a class of ligand-activated transcription factors that regulate diverse biological processes including development, homeostasis, and metabolism [1, 2]. Like other nuclear receptors, AR is composed of several modular domains, including a DNA-binding domain, a ligand-binding domain, and an intrinsically flexible N-terminal domain that mediates transcriptional regulation. Upon binding androgenic hormones such as testosterone or dihydrotestosterone (DHT), AR undergoes conformational rearrangements that promote dissociation from cytoplasmic chaperone complexes and enable its translocation into the nucleus [3, 4]. Within the nucleus, ligand-bound AR dimerizes and binds androgen response elements (AREs), recruiting transcriptional coactivators that regulate gene expression [5, 6].

Androgenic hormones are steroidal molecules synthesized primarily in the testes and adrenal glands and are essential for male sexual development and reproductive function [7]. Dysregulation of AR signaling has been implicated in a wide range of pathologies. In particular, aberrant AR activity is a defining feature of prostate cancer progression, including castration-resistant prostate cancer (CRPC), where AR signaling persists despite androgen deprivation [8, 9]. Conversely, loss-of-function mutations in AR lead to androgen insensitivity syndrome, underscoring the receptor’s central physiological role [10]. In addition to its canonical genomic functions, AR also participates in rapid non-genomic signaling pathways through interactions with cytoplasmic or membrane-associated proteins, influencing cellular proliferation, survival, and migration [11].

Recent work by Bajpai et al. [12] revealed an unexpected aspect of AR biology: its ability to localize to mitochondria in vivo. Deletion of the first 36 residues of the AR N-terminus abolished mitochondrial import, indicating that this region contains a functional mitochondrial localization sequence (MLS). Fusion of this N-terminal segment to green fluorescent protein (GFP) resulted in robust mitochondrial localization, demonstrating that the AR N-terminus is sufficient to direct a heterologous protein to mitochondria [13]. GFP-based reporter assays provide a powerful means of visualizing subcellular targeting, as GFP fluorescence allows direct observation of mitochondrial localization when co-localized with organelle-specific markers [14].

Computational analysis using MitoProt further supported these experimental findings, predicting a high probability of mitochondrial targeting and identifying a potential cleavage site within the first 14 residues of the AR sequence [15]. Together, these results establish the AR N-terminal region as a bona fide mitochondrial targeting signal and motivate a detailed investigation of its underlying structural properties.

Guided by this evidence, we isolated the first 15 residues of the AR sequence (MEVQLGLGRVYPRPP), hereafter referred to as the mitochondrial localization peptide (MLP), and examined its conformational behavior. The selection of this sequence encompasses the predicted MLS and cleavage site while allowing for potential flanking effects at the C-terminus [12]. Notably, MLP is intrinsically disordered, lacking a stable tertiary structure under physiological conditions. Intrinsically disordered peptides and proteins (IDPs) exist as dynamic ensembles of rapidly interconverting conformations rather than adopting a single well-defined fold [16, 17, 18, 19]. This conformational plasticity is often essential for biological function, particularly in processes requiring transient interactions, molecular recognition, and cellular targeting [20].

IDPs play central roles in regulatory networks, enabling flexible and tunable responses to cellular signals through mechanisms such as conformational selection and binding-induced folding [21]. At the same time, intrinsic disorder is strongly associated with disease-related proteins, emphasizing the importance of understanding how sequence features shape disordered conformational ensembles [22]. Mitochondrial targeting peptides, which are typically short, enriched in amphipathic and flexible residues, and lack persistent secondary structure, represent a particularly compelling class of IDPs whose function is intimately linked to dynamic structural behavior.

In this study, we systematically investigate the conformational dynamics of the wildtype MLP and a complete panel of 19 single-point variants generated by substitution of the second residue, originally a glutamic acid. The second position was selected due to its location within the predicted MLS, where electrostatic and structural properties are expected to strongly influence targeting behavior [12]. By generating multiple independent replicas for each variant, we capture the inherent heterogeneity of the disordered ensemble and enable statistically robust comparisons across sequences.

Using extensive all-atom molecular dynamics simulations, we characterize how single-residue substitutions subtly reshape both global and local features of the MLP conformational ensemble. We analyze compactness, residue-level secondary structure propensities, and higher-order similarities among variants to identify mutations that preserve wild-type-like behavior and those that produce more pronounced deviations. To further probe the limits of equilibrium sampling for intrinsically disordered systems, we apply enhanced sampling approaches to selected variants and assess their ability to converge the underlying free-energy landscape.

Together, this work provides a systematic and quantitative framework for linking minimal sequence variations to changes in the dynamic structural ensemble of a mitochondrial localization peptide. More broadly, it highlights both the sensitivity of disordered targeting signals to local sequence context and the methodological challenges inherent in fully characterizing intrinsically disordered peptides.

## Methods

### Peptide System Preparation

A model 15-residue peptide (MEVQLGLGRVYPRPP) was designed, corresponding to the mitochondrial localization sequence (MLS) identified within the N-terminal domain of the androgen receptor (AR) by Bajpai et al. [12]. This sequence, referred to hereafter as the wild-type mitochondrial localization peptide (MLP), encompasses the residues predicted to contain the critical targeting information necessary for mitochondrial import.

To systematically probe the impact of sequence variation on peptide conformational dynamics, we generated a full panel of 20 variants by substituting the glutamic acid at the second position (E2) with each of the other 19 standard amino acids, resulting in sequences labeled MLP_wt_, MLP_E2A_, MLP_E2C_, …, MLP_E2Y_.

Initial structures for each peptide variant were generated using the LEaP module from the AmberTools suite [23]. Linear (extended) conformations were constructed from the amino acid sequences and subsequently subjected to energy minimization to relieve steric clashes. To capture the conformational heterogeneity typical of intrinsically disordered peptides (IDPs), 16 distinct starting conformations were generated for each sequence through random perturbations and short equilibration simulations.

All systems were prepared using a generalized Born implicit solvent (GBIS) model [24, 25] to approximate solvation effects while enabling faster conformational sampling compared to explicit solvent simulations. This resulted in a comprehensive dataset comprising 320 independent peptide systems (20 variants × 16 initial conformations) for subsequent molecular dynamics simulations and analysis.

### Molecular Dynamics Simulations

All equilibrium molecular dynamics simulations were performed using NAMD 2.14 [26], a highly parallelized MD engine optimized for large-scale biomolecular systems. Simulations employed either the CHARMM36 [27] or the updated CHARMM36m [28] force field, depending on the specific system under study. These force fields have been extensively validated for accurate modeling of both folded and intrinsically disordered proteins.

Each peptide system was modeled at a physiological temperature of 310 K and a salt concentration of 0.15 M NaCl. Solvent effects were treated using the generalized Born implicit solvent (GBIS) model [24, 25], which enables faster sampling compared to explicit solvent representations while maintaining reasonable accuracy for small peptides.

Each initial structure underwent a 250 ps equilibration phase to relax any unfavorable contacts, followed by a 100 ns production run. For each peptide variant, 16 independent simulations were conducted, resulting in a total sampling time of 1.6 *μ*s per variant. This simulation setup provided a robust statistical basis for analyzing the conformational dynamics and secondary structure propensities of the mitochondrial localization peptide (MLP) variants.

### Enhanced Sampling Simulations

To achieve deeper sampling of the conformational landscape beyond equilibrium simulations, a subset of four peptide variants—MLP_wt_, MLP_E2A_, MLP_E2K_, and MLP_E2Q_—was subjected to enhanced sampling via well-tempered metadynamics (WTM) [29].

All metadynamics simulations were performed using NAMD 2.14 [26], starting from representative structures selected from the equilibrium GBIS trajectories. The *α*-helical content was employed as the collective variable (CV) to bias the system away from previously visited conformational states and encourage exploration of alternative structural ensembles.

Two WTM simulation protocols were employed sequentially:

- **Coarse WTM**: Simulations were carried out with a Gaussian hill width of 0.05 and a hill height of 0.5, over a simulation timespan of 0.5 *μ*s per system.
- **Fine WTM**: Subsequently, finer-grained simulations were performed with a hill width of 0.01 and a hill height of 0.1, extending the simulation time to 2.5 *μ*s per system.

Initially, metadynamics simulations were conducted under the generalized Born implicit solvent (GBIS) model [24], but were discontinued after insufficient convergence was observed. To improve sampling fidelity, explicit solvent systems were constructed using the psfgen module from the Visual Molecular Dynamics (VMD) package [30], solvated in a cubic box of TIP3P water molecules [31], and ionized to 0.15 M NaCl. These explicit systems underwent energy minimization, followed by a 2.5 ns equilibration run prior to initiating fine WTM simulations.

This enhanced sampling approach enabled broader exploration of peptide conformational space, necessary for capturing the diverse structural fluctuations inherent to intrinsically disordered peptides.

### Analysis

To characterize the conformational behavior of the mitochondrial localization peptide (MLP) variants, a comprehensive suite of structural analyses was conducted on the equilibrium MD trajectories.

Basic compactness metrics were first computed for each system. The radius of gyration (*R*_*g*_), end-to-end distance, and solvent accessible surface area (SASA) were calculated using built-in plugins in Visual Molecular Dynamics (VMD) [30]. These parameters provided a coarse-grained assessment of conformational compactness and overall molecular extension. All measurements were averaged across the 16 replicas for each variant to obtain robust ensemble statistics.

To further dissect residue-level structural preferences, backbone dihedral angles (*ϕ, ψ*) were extracted from each frame of the simulation trajectories. Each sampled conformation was categorized into one of five secondary structure motifs—right-handed *α*-helix (*α*R), left-handed *α*-helix (*α*L), polyproline II helix (PPII), *β*-sheet, or unassigned—based on the Ramachandran region classification described by Moradi et al. [32]. For each residue, the occupancy (percentage of simulation time spent) in each structural category was calculated and averaged across replicas, enabling comparison of secondary structure propensities across different mutants.

Given the large dataset generated (320 trajectories across 20 peptide variants), full Ramachandran plots were not plotted for each system. Instead, secondary structural propensity profiles were extracted to enable aggregate, residue-by-residue comparisons without sacrificing interpretability.

To reduce the complexity of the resulting high-dimensional dataset, principal component analysis (PCA) was employed. For each trajectory, the average occupancy of three broad secondary structure classes (right-handed *α*-helix, combined *β*-sheet/PPII, and unstructured) for each residue was used as the input feature vector. The dataset, consisting of 320 individual trajectories and 20 averaged profiles, was subjected to PCA using standard algorithms [33]. This dimensionality reduction technique enabled visualization of overall trends and clustering among mutants based on their secondary structural behaviors.

Mutant ranking was performed by calculating the relative distance of each variant from the wild-type MLP in the reduced principal component space. Variants such as MLP_E2H_, MLP_E2I_, MLP_E2M_, and MLP_E2W_ clustered closely with wild type, suggesting minimal perturbation of overall structural dynamics. In contrast, variants like MLP_E2L_ and MLP_E2P_ showed greater divergence. Visualization strategies employed color coding, where red markers indicated sequences validated by experimental data, and green markers denoted novel predictions warranting further exploration.

## Results

### Global Compactness and Baseline Conformational Properties of MLP Variants

To establish a baseline description of the conformational behavior of the mitochondrial localization peptide (MLP), we first examined global compactness metrics across selected sequence variants using equilibrium molecular dynamics simulations. These analyses were intended to assess whether mutations at the second residue position induce large-scale changes in peptide dimensions or solvent exposure, prior to more detailed residue-level characterization.

Representative global metrics for the wild-type peptide and three initial variants (E2A, E2K, and E2Q) are summarized in Table 1. Across these systems, the average radius of gyration, end-to-end distance, and solvent-accessible surface area (SASA) differ only modestly, with variations well within the spread expected for intrinsically disordered peptides sampled across independent trajectories. None of the substitutions examined produce a pronounced shift toward either significantly more compact or more extended conformational ensembles, indicating that mutation at position 2 does not strongly perturb overall chain dimensions at this level of analysis.

**Table 1:** Preliminary compactness metrics for selected MLP variants. Each value represents an average across 16 independent parallel simulations.

| Variant | Radius of Gyration ($\text{\AA}$) | End-to-End Distance ($\text{\AA}$) | SASA ($\text{\AA}^2$) |
| --- | --- | --- | --- |
| MLP<sub>wt</sub> | 9.88 | 19.05 | 1576.14 |
| MLP<sub>E2A</sub> | 9.89 | 19.20 | 1580.63 |
| MLP<sub>E2K</sub> | 9.94 | 19.24 | 1583.09 |
| MLP<sub>E2Q</sub> | 9.92 | 19.25 | 1583.28 |

Consistent with these observations, visual inspection of representative conformations sampled during equilibrium simulations (Figure 1) reveals broadly heterogeneous ensembles lacking persistent tertiary structure. All variants populate a wide range of extended and partially collapsed conformations, characteristic of intrinsically disordered systems, with no single dominant fold emerging over the course of the simulations.

**Figure 1:**
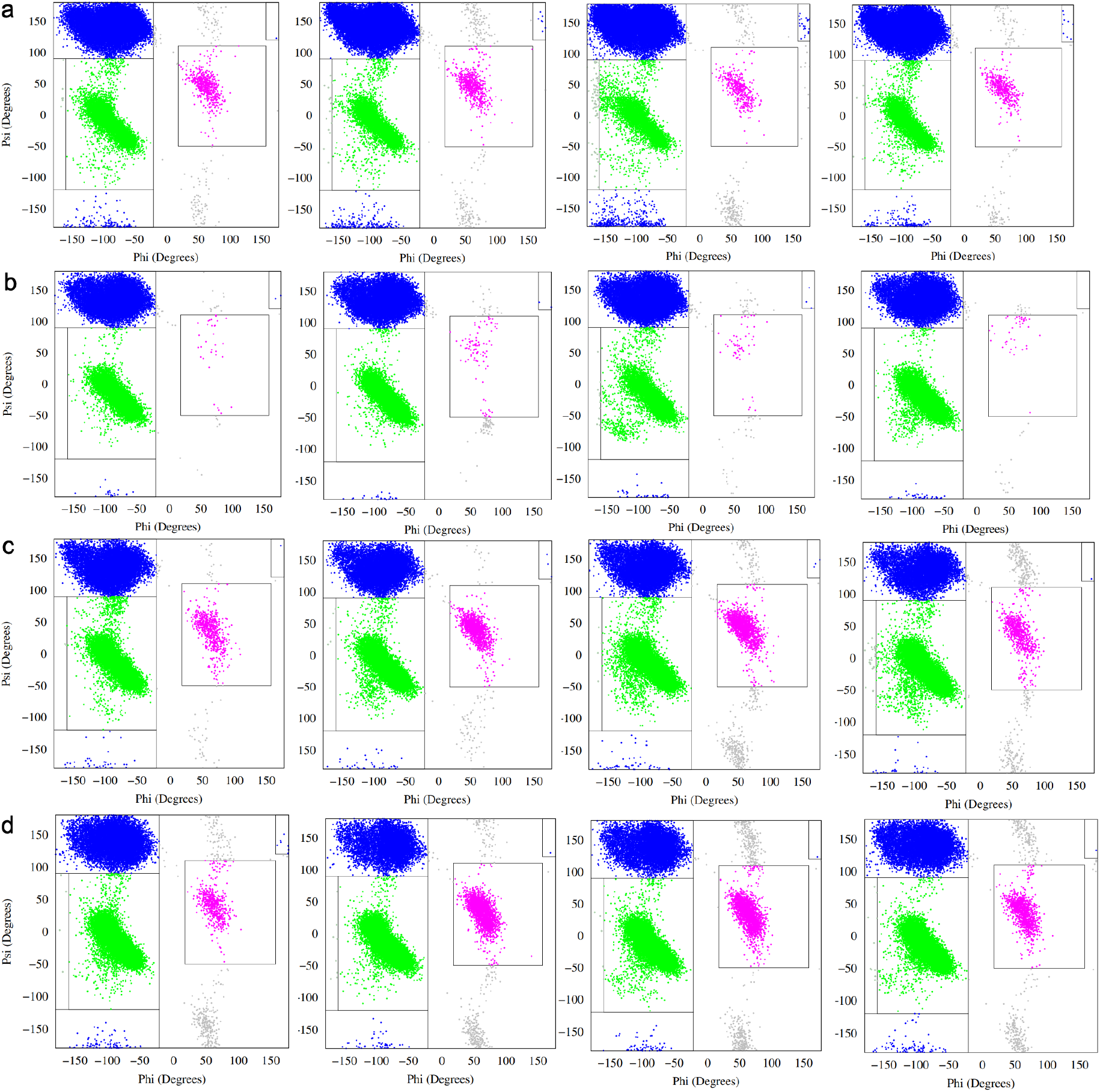
a) Ramachandran plot denoting secondary structural propensity for MLP-wt, MLP-E2A, MLP-E2K and MLP-E2Q (left to right) (residue 2), variable b) Ramachandran secondary plot for MLP-wt, MLP-E2A, MLP-E2K and MLP-E2Q (left to right) (residue 3), valine c) Ramachandran secondary plot for MLP-wt, MLP-E2A, MLP-E2K and MLP-E2Q (left to right) (residue 4), glutamine d) Ramachandran secondary plot for MLP-wt (residue 5), leucine

These results indicate that global compactness metrics alone are insufficient to distinguish between MLP variants, despite the chemical diversity of the substituted residues. This lack of strong separation suggests that any mutation-induced effects are likely subtle and localized, manifesting primarily through shifts in transient secondary structural preferences rather than through large-scale changes in peptide size or solvent exposure. This conclusion motivates the more detailed residue-resolved and ensemble-based analyses presented in subsequent sections.

Importantly, the limited discriminatory power of global metrics underscores a central challenge in the computational characterization of intrinsically disordered peptides: ensemble-averaged scalar observables such as radius of gyration or SASA may mask meaningful differences in underlying conformational populations. As such, more sensitive descriptors—capable of resolving local structural propensities and correlated dynamical behavior—are required to capture the effects of sequence variation within the MLP ensemble.

### Residue-Resolved Secondary Structural Propensities Revealed by Ramachandran Analysis

Given the limited ability of global compactness metrics to discriminate between MLP variants, we next examined residue-level conformational behavior by analyzing backbone dihedral angle sampling across the peptide sequence. For each variant, the *ϕ/ψ* angles of all residues were monitored throughout the equilibrium trajectories and mapped onto Ramachandran space, allowing transient secondary structural propensities to be quantified on a per-residue basis.

Representative Ramachandran plots for selected residues near the N-terminus (residues 2–5) are shown for the wild-type peptide and three initial variants (E2A, E2K, and E2Q) in Figure 2. Across all systems, the distributions are broad and diffuse, consistent with intrinsically disordered behavior and the absence of stable secondary structure. Sampling is dominated by regions corresponding to *β* and polyproline II (PPII) conformations, with intermittent population of right-handed and left-handed *α*-helical regions. No residue exhibits persistent localization to a single Ramachandran basin over the course of the simulations.

**Figure 2:**
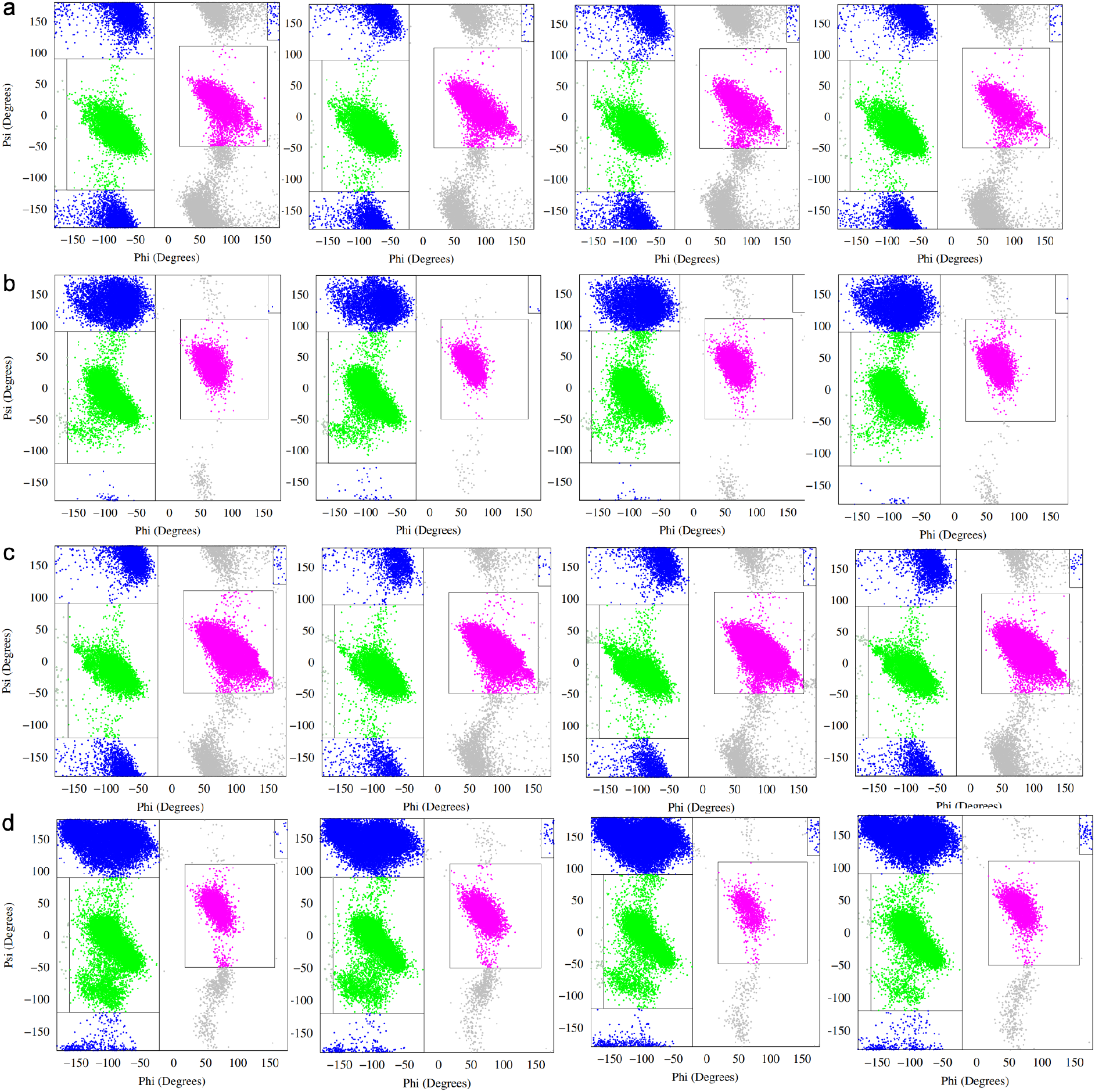
a) Ramachandran plot denoting secondary structural propensity for MLP-wt, MLP-E2A, MLP-E2K and MLP-E2Q (left to right) (residue 6), glycine b) Ramachandran secondary plot for MLP-wt, MLP-E2A, MLP-E2K and MLP-E2Q (left to right) (residue 7), leucine c) Ramachandran secondary plot for MLP-wt, MLP-E2A, MLP-E2K and MLP-E2Q (left to right) (residue 8), glycine d) Ramachandran secondary plot for MLP-wt (residue 9), arginine

**Figure 3:**
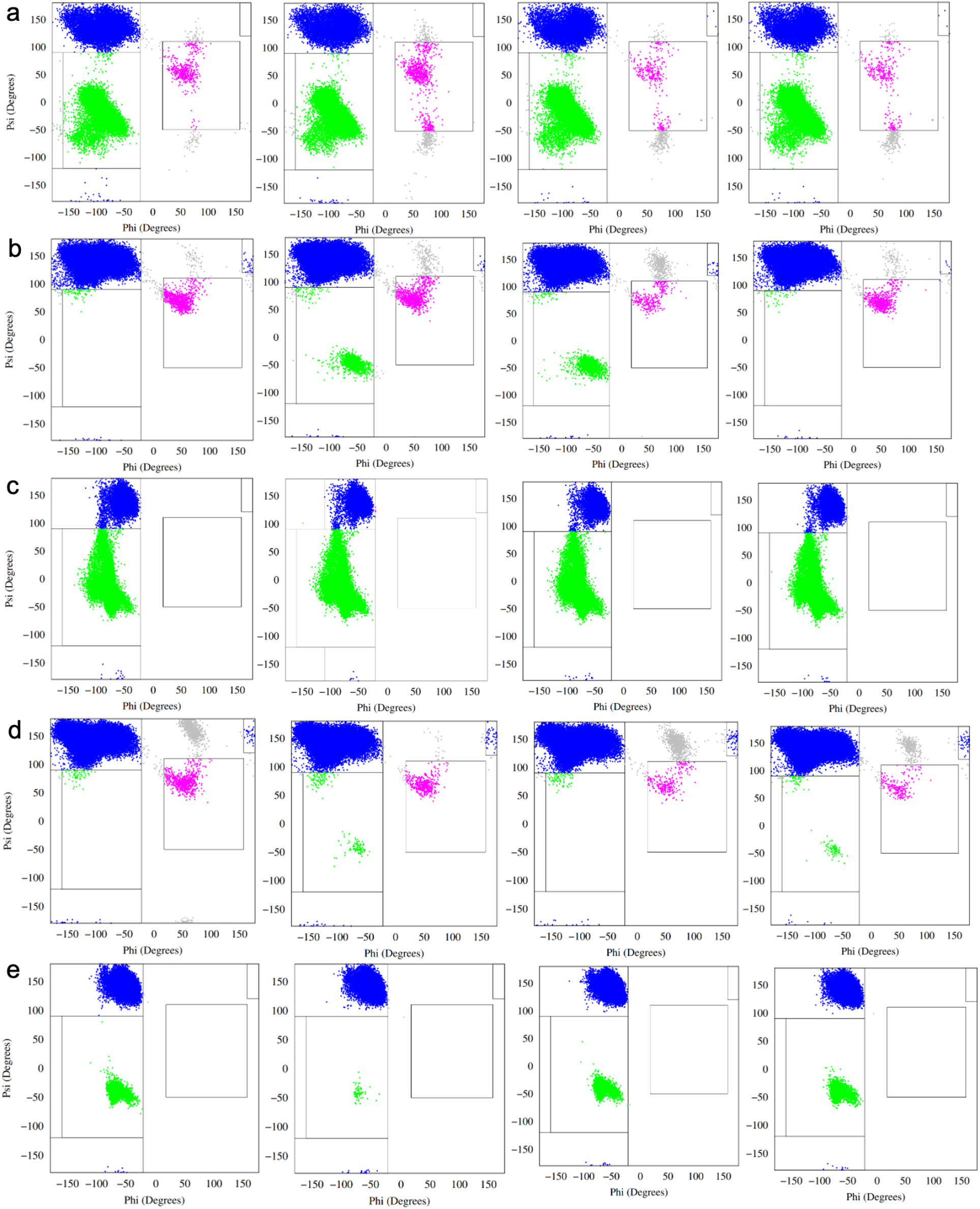
a) Ramachandran plot denoting secondary structural propensity for MLP-wt, MLP-E2A, MLP-E2K and MLP-E2Q (left to right) (residue 10), valine b) Ramachandran secondary plot for MLP-wt, MLP-E2A, MLP-E2K and MLP-E2Q (left to right) (residue 11), tyrosine c) Ramachandran secondary plot for MLP-wt, MLP-E2A, MLP-E2K and MLP-E2Q (left to right) (residue 12), proline d) Ramachandran secondary plot for MLP-wt (residue 13), arginine e) Ramachandran secondary plot for MLP-wt (residue 14), proline

Despite this overall disorder, subtle but reproducible shifts in sampling are evident upon mutation at the second residue position. In particular, differences in the relative occupancy of *α*-helical versus *β*/PPII regions are observed not only at the mutated site itself, but also at neighboring residues downstream. This indicates that substitutions at position 2 can modulate local backbone conformational preferences beyond their immediate location, likely through changes in local electrostatics, sterics, or backbone flexibility.

To quantify these effects more systematically, Ramachandran populations were classified into discrete secondary structural categories, and the resulting occupancies were averaged across the 16 independent simulations for each variant. Deviations in secondary structural propensity relative to the wild-type ensemble are summarized for E2A, E2K, and E2Q in Figure 4. These plots reveal that mutations at position 2 produce variant-specific shifts in local secondary structure that are distributed unevenly along the sequence. The most pronounced deviations occur near the N-terminal region and around glycine-rich segments, whereas residues within the proline-rich C-terminal region show comparatively smaller changes, consistent with their strong intrinsic bias toward PPII-like conformations.

**Figure 4:**
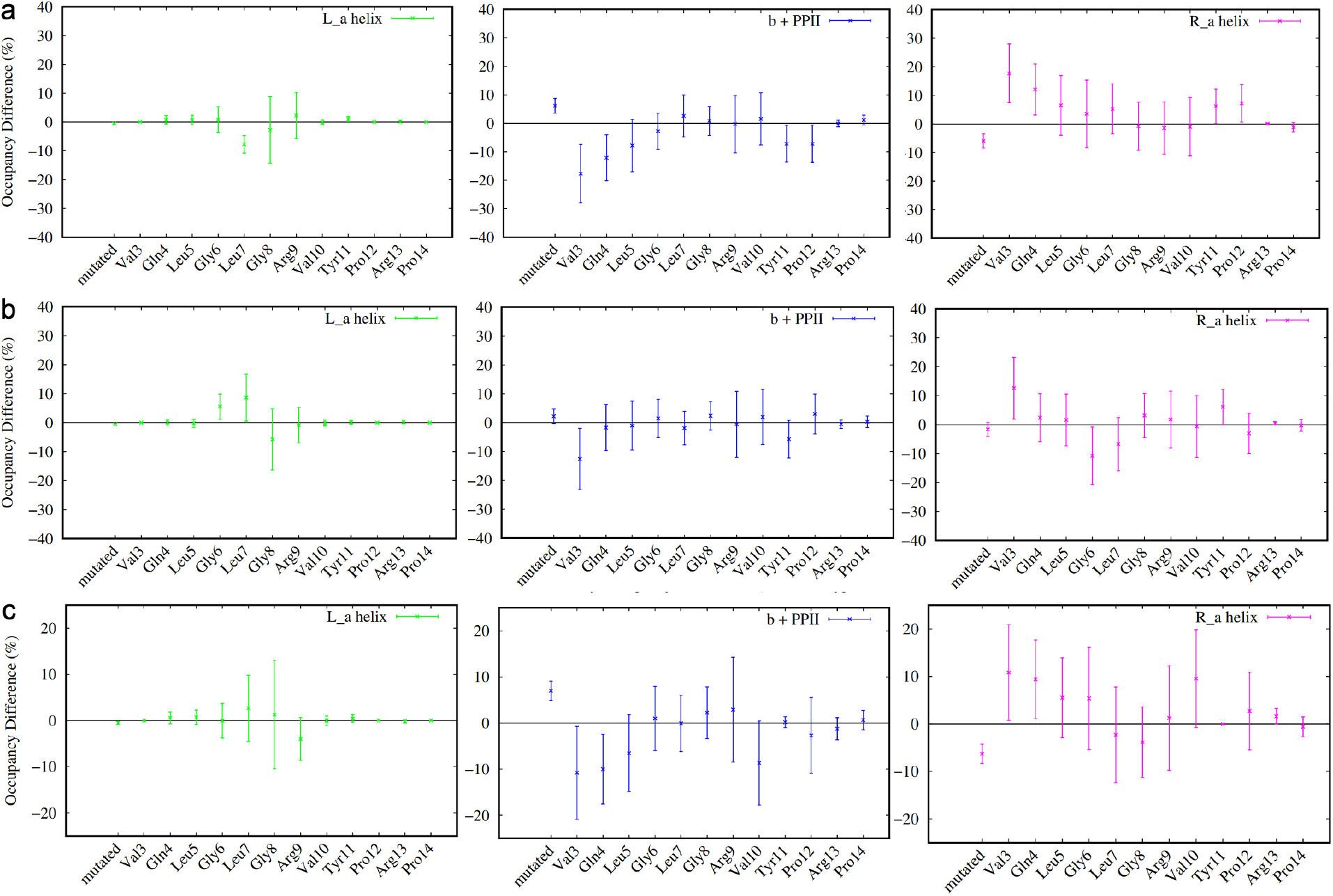
Sample plots of secondary structural deviance of a) mlp e2a vs wild type, b) mlp e2k vs wild type and c) mlp e2q vs wild type. Occupancies are derived from Ramachandran plots as a ratio of how much time a given residue spends in each region and reflect an average of the 16 parallel simulations. Error bars reflect the appropriate spread of averages from both data sets.

Importantly, the magnitude of these deviations is modest, and error bars across parallel simulations remain substantial, reflecting the inherent variability of intrinsically disordered ensembles. This variability underscores the challenge of extracting statistically robust conclusions from individual residues in isolation. Nevertheless, the consistency of the observed trends across independent simulations suggests that the detected differences reflect genuine ensemble-level shifts rather than stochastic noise.

The full set of residue-resolved deviation plots for all sequence variants is provided in Figures S1–S7. Inspection of these data reveals that certain substitutions produce secondary structural patterns that more closely resemble the wild-type ensemble, while others induce more pronounced redistribution of conformational sampling across multiple residues. Notably, these effects are not uniformly distributed across the peptide, indicating that the influence of the second residue position propagates through the sequence in a residue- and context-dependent manner.

Taken together, these results demonstrate that while MLP variants are globally similar in size and disorder, mutation at a single position can measurably reshape the local conformational landscape of the peptide. However, because these changes are subtle and distributed across many residues, a higher-level, multivariate description of ensemble behavior is required to compare variants in a systematic and unbiased manner. This motivates the dimensionality-reduction and clustering analysis presented in the following section.

### Multivariate Classification of MLP Variants Using Principal Component Analysis

While residue-resolved secondary structural analyses reveal subtle but reproducible differences between MLP variants, the distributed and correlated nature of these changes makes direct comparison challenging when considered on a residue-by-residue basis. To provide a higher-level, unbiased description of ensemble behavior across all variants, we applied principal component analysis (PCA) to the secondary structural propensity data derived from the Ramachandran classification.

For this analysis, each simulation was treated as an independent observation, with variables defined by the average occupancies of discrete secondary structural states for each residue along the peptide. This representation captures the collective conformational tendencies of the peptide while reducing sensitivity to instantaneous fluctuations inherent to intrinsically disordered systems. PCA was then used to project this high-dimensional dataset onto a reduced set of orthogonal components that capture the dominant sources of variance across all simulations.

Figure 5 shows the projection of individual simulations for the wild-type peptide and representative variants onto the first two principal components. In these plots, each point corresponds to a single equilibrium trajectory, while the mean behavior of each variant is indicated separately. The broad dispersion of individual trajectories within each variant reflects the substantial conformational heterogeneity of the MLP ensemble, consistent with its intrinsically disordered nature. Notably, the spread of wild-type trajectories spans a large region of the projected space, highlighting the difficulty of defining a single representative structure or narrow conformational basin for this system.

**Figure 5:**
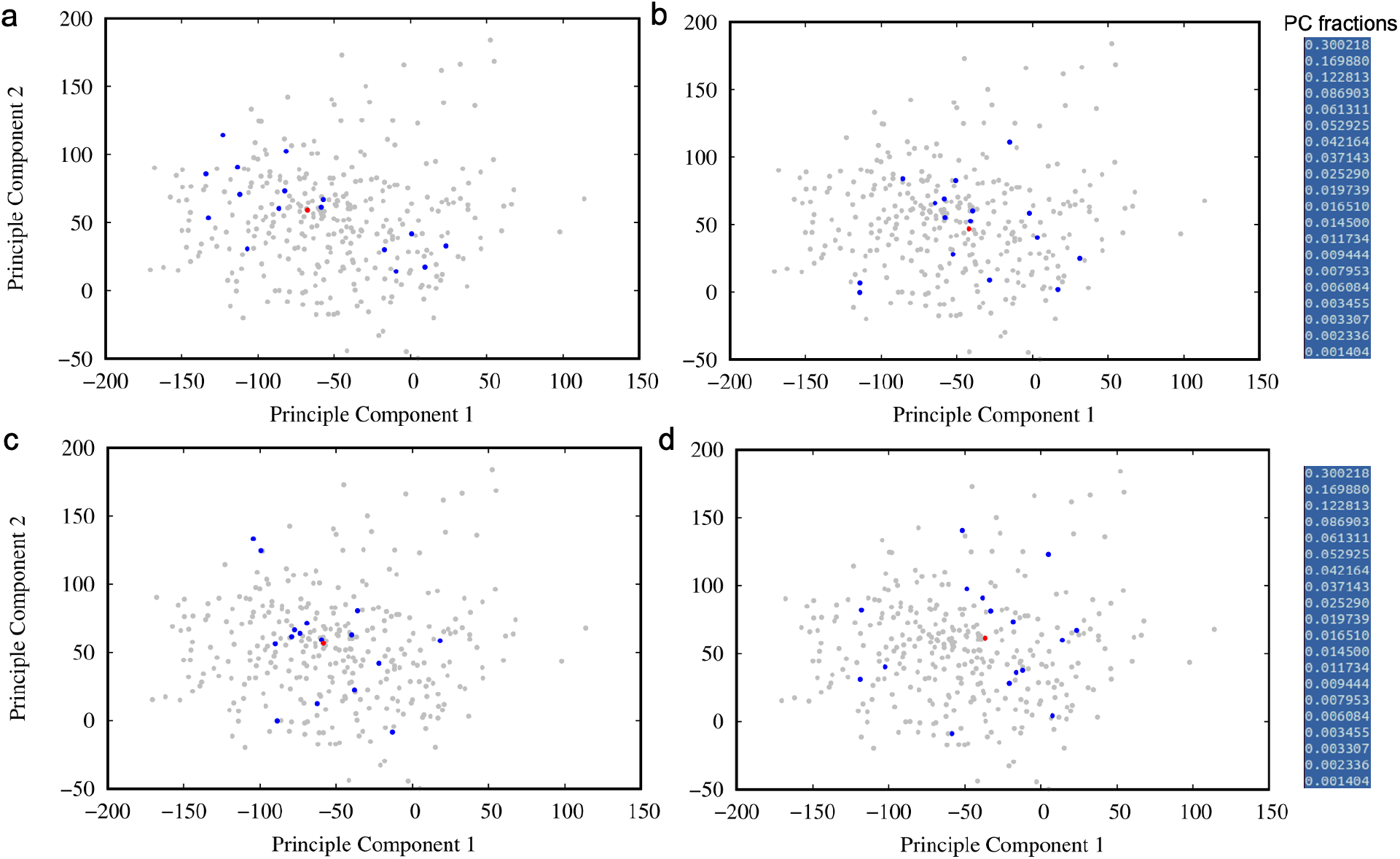
Principal component projection of secondary structural propensity data of a) mlp e2a, b) mlp e2k, c) mlp e2q, and d) mlp wild type. Each blue dot represents one of 16 parallel simulations, while red dot represents the averaged behavior over all 16 simulations. Grey dots show the spread of all remaining variants.

Despite this heterogeneity, systematic differences between variants emerge when considering the relative positions of their ensemble-averaged behaviors. The clustering of variant means in PCA space suggests that certain substitutions at position 2 give rise to secondary structural ensembles that are more similar to the wild type, while others exhibit more pronounced deviations. These relationships are more clearly visualized when considering only the averaged behavior of each variant, as shown in Figures S8–S10.

Inspection of the PCA projections reveals that variants with similar physicochemical properties at the second position tend to occupy nearby regions of the reduced-dimensional space, indicating that substitution chemistry can bias the global conformational ensemble in a reproducible manner. Conversely, several variants appear displaced from the wild-type region, reflecting more substantial redistribution of secondary structural propensities across the sequence. Importantly, these distinctions arise from collective, multiresidue effects rather than from localized changes at a single position, underscoring the value of a multivariate description of ensemble behavior.

It is important to emphasize that the PCA results do not imply sharp boundaries or discrete conformational states separating variants. Instead, the observed clustering should be interpreted as a probabilistic tendency for certain sequences to sample overlapping or distinct regions of conformational space. The loose clustering and partial overlap between variants further indicate that extensive sampling is required to fully characterize these ensembles and that conclusions drawn from individual trajectories should be made cautiously.

Nevertheless, the PCA framework provides a useful means of organizing and comparing the complex secondary structural behavior of MLP variants in a systematic manner. By reducing a high-dimensional, residue-resolved dataset into a small number of collective descriptors, this approach enables qualitative classification of variants according to their overall conformational tendencies. These classifications, in turn, inform subsequent analyses aimed at assessing sampling limitations and exploring enhanced sampling strategies for more complete characterization of the MLP conformational landscape.

### Enhanced Sampling Highlights Persistent Ruggedness of the MLP Conformational Landscape

The equilibrium simulations and multivariate analyses presented above indicate that MLP variants populate broad and heterogeneous conformational ensembles, with subtle sequence-dependent biases that are difficult to resolve conclusively within accessible equilibrium timescales. To further assess the extent of conformational sampling and to probe the underlying freeenergy landscape, we performed enhanced sampling simulations using well-tempered metadynamics (WTM) for a subset of representative variants.

In these simulations, the collective variable was chosen to reflect the overall *α*-helical content of the peptide, with the goal of accelerating transitions between conformations differing in transient secondary structure. WTM simulations were carried out in explicit solvent for the wild-type peptide and three initial variants (E2A, E2K, and E2Q), extending to multi-microsecond timescales. Free-energy profiles were reconstructed from the accumulated bias at successive fractions of the trajectory in order to evaluate convergence behavior and landscape stability.

Time-resolved potentials of mean force (PMFs) for all four variants are shown in Figures S11–S14. Across systems, the PMFs exhibit pronounced evolution as additional bias is deposited, with changes persisting into the final portions of the trajectories. In particular, while low-helicity regions remain consistently favorable and highly helical states are generally disfavored, the intermediate helicity regime displays substantial reshaping over time, including the appearance, disappearance, or shifting of metastable features.

The continued evolution of these PMFs indicates that convergence is not fully achieved even after several microseconds of enhanced sampling. This behavior suggests that *α*-helical content alone does not constitute a sufficient collective variable to capture all of the slow degrees of freedom governing MLP conformational dynamics. Orthogonal motions, such as backbone bending, glycine- and proline-mediated flexibility, or transitions between *β* and polyproline II conformations, likely contribute significantly to the overall free-energy landscape but are not directly accelerated by the chosen bias.

Importantly, these observations are consistent across all examined variants, indicating that the lack of convergence is not specific to a particular sequence but rather reflects the intrinsic complexity of the MLP conformational ensemble. As such, the metadynamics results should be interpreted as diagnostic rather than quantitative, illustrating both the ruggedness of the underlying landscape and the limitations of single-variable biasing schemes for intrinsically disordered peptides.

Taken together, the enhanced sampling simulations reinforce the conclusions drawn from equilibrium and PCA-based analyses: MLP exhibits a highly heterogeneous and multidimensional conformational landscape that is sensitive to sequence variation yet difficult to exhaustively sample. These findings motivate the use of more sophisticated enhanced sampling strategies, such as bias-exchange or multi-dimensional biasing approaches, to achieve more complete characterization of the conformational space and to enable more definitive comparisons between sequence variants.

## Discussion

In this work, we used extensive equilibrium molecular dynamics simulations combined with ensemble-based analysis to investigate the conformational behavior of an intrinsically disordered mitochondrial localization peptide (MLP) and a systematic set of sequence variants. By focusing on mutations at a single, chemically diverse position near the N-terminus, we sought to assess how subtle sequence changes modulate conformational ensembles in the absence of stable tertiary structure.

A central finding of this study is that mutation at position 2 does not produce large-scale changes in global peptide dimensions. Across examined variants, measures such as radius of gyration, end-to-end distance, and solvent-accessible surface area remain broadly similar, consistent with highly disordered ensembles. These observations underscore a key limitation of global scalar descriptors when applied to intrinsically disordered systems: ensemble-averaged measures of size or compactness may obscure meaningful differences in underlying conformational populations.

In contrast, residue-resolved analyses reveal that substitutions at position 2 induce subtle but reproducible shifts in local secondary structural propensities. Ramachandran-based classification shows that these effects extend beyond the mutated residue itself, influencing neighboring regions along the peptide backbone. In particular, changes in the balance between *α*-helical, *β*, and polyproline II conformations are most evident near the N-terminus and within glycine-rich segments, whereas the proline-rich C-terminal region exhibits comparatively robust conformational bias. These observations suggest that even in short intrinsically disordered peptides, local sequence context and backbone flexibility can mediate the propagation of conformational effects over multiple residues.

Because these sequence-dependent shifts are distributed across the peptide and often small in magnitude, direct comparison of variants based on individual residues is challenging. To address this, we employed principal component analysis to construct a multivariate representation of secondary structural behavior across the full ensemble. PCA reveals that while individual trajectories exhibit substantial dispersion, ensemble-averaged behaviors cluster in a sequence-dependent manner. Certain substitutions yield conformational tendencies closely resembling the wild-type ensemble, whereas others produce more pronounced deviations. Importantly, these distinctions emerge only when secondary structural information is considered collectively, reinforcing the value of ensemble-level and multivariate approaches for characterizing intrinsically disordered systems.

Enhanced sampling simulations further illuminate the complexity of the MLP conformational landscape. Well-tempered metadynamics biasing the overall *α*-helical content accelerates exploration of conformations differing in transient helicity but does not yield fully converged free-energy surfaces, even on multi-microsecond timescales. Persistent evolution of the reconstructed potentials of mean force, particularly within intermediate helicity regimes, indicates that *α*-helical content alone is insufficient to capture all slow collective motions governing MLP dynamics. These findings highlight the inherently multidimensional nature of the conformational landscape and emphasize the limitations of single-variable biasing schemes when applied to intrinsically disordered peptides.

Taken together, the results of this study illustrate both the sensitivity and the challenges associated with computational characterization of intrinsically disordered localization sequences. While sequence variation at a single position can measurably bias local and global conformational tendencies, these effects manifest as shifts in ensemble probabilities rather than as discrete structural transitions. Consequently, rigorous interpretation requires careful consideration of sampling limitations, ensemble variability, and the choice of structural descriptors.

From a biological perspective, the observed modulation of transient secondary structural propensities near the N-terminus is noteworthy in the context of mitochondrial targeting. Mitochondrial localization peptides are often characterized by transient amphipathic helices that are stabilized upon interaction with import machinery. Although the present simulations do not address binding or translocation directly, the sequence-dependent shifts in helicity observed here may influence the likelihood of sampling such conformations, thereby providing a potential mechanistic link between sequence composition and localization behavior. Future studies integrating explicit interactions with mitochondrial membranes or import receptors will be required to evaluate this hypothesis directly.

More broadly, this work demonstrates a generalizable computational framework for dissecting the conformational consequences of sequence variation in intrinsically disordered peptides. By combining parallel equilibrium simulations, residue-resolved secondary structure analysis, and multivariate classification, it is possible to extract meaningful ensemble-level trends even in the absence of converged free-energy landscapes. The enhanced sampling results further motivate the development and application of higher-dimensional biasing strategies, such as bias-exchange or dihedral-based metadynamics, to more fully resolve the complex conformational landscapes of disordered targeting sequences.

## Conclusion

In this study, we employed extensive molecular dynamics simulations and ensemble-based analysis to investigate the conformational behavior of an intrinsically disordered mitochondrial localization peptide and a systematic set of sequence variants. By focusing on substitutions at a single residue position near the N-terminus, we examined how localized sequence changes influence conformational sampling in the absence of stable secondary or tertiary structure.

Our results demonstrate that global measures of compactness, such as radius of gyration, end-to-end distance, and solvent-accessible surface area, are largely insensitive to sequence variation within the MLP ensemble. While these metrics confirm the highly disordered nature of the peptide, they do not adequately capture mutation-dependent differences in conformational behavior. In contrast, residue-resolved analysis of backbone dihedral sampling reveals subtle but reproducible shifts in secondary structural propensities that extend beyond the mutated site and propagate along the peptide backbone in a sequence-dependent manner.

By integrating these local structural descriptors into a multivariate framework, principal component analysis enables systematic comparison of MLP variants at the ensemble level. Although individual simulations exhibit substantial variability, ensemble-averaged behaviors cluster in a manner that reflects underlying sequence chemistry, illustrating how collective analysis can reveal trends that are not apparent from single observables or individual trajectories. These findings underscore the importance of ensemble-level and multidimensional descriptors for characterizing intrinsically disordered systems.

Enhanced sampling simulations further highlight the rugged and high-dimensional nature of the MLP conformational landscape. Well-tempered metadynamics biasing *α*-helical content accelerates exploration of transient secondary structure but does not achieve full convergence on accessible timescales, emphasizing the limitations of single-variable biasing strategies for intrinsically disordered peptides. Together with the equilibrium results, these observations point to the need for more sophisticated enhanced sampling approaches to fully resolve the conformational space of disordered localization sequences.

Taken together, this work provides a detailed and methodologically rigorous characterization of how minimal sequence variation can bias the conformational ensembles of an intrinsically disordered targeting peptide. Beyond the specific system studied here, the analytical framework developed in this work is broadly applicable to other intrinsically disordered peptides and localization sequences, offering a general strategy for extracting meaningful structure–function insights from heterogeneous and highly dynamic molecular ensembles.

## Acknowledgments

This research was supported by the National Institute of General Medical Sciences (NIH grant R35GM147423 awarded to M.M.), the National Science Foundation (NSF grant CHE 1945465 awarded to M.M.), and the Arkansas Biosciences Institute. A.F. acknowledges Emad Tajkhorshid for his continuous support and insightful discussions and for the funding through the NIH grant R24-GM145965. The work used Stampede at the Texas Advanced Computing Center and Bridges-2 at the Pittsburgh Supercomputing Center through allocation MCB150129 to M.M. from the Advanced Cyberinfrastructure Coordination Ecosystem: Services & Support (ACCESS) program. Additional computational support came from the Arkansas High-Performance Computing Center, funded by multiple NSF grants and the Arkansas Economic Development Commission.

