## Supporting Information for "Characterizing the Conformational Dynamics of an Intrinsically Disordered Localization Sequence"

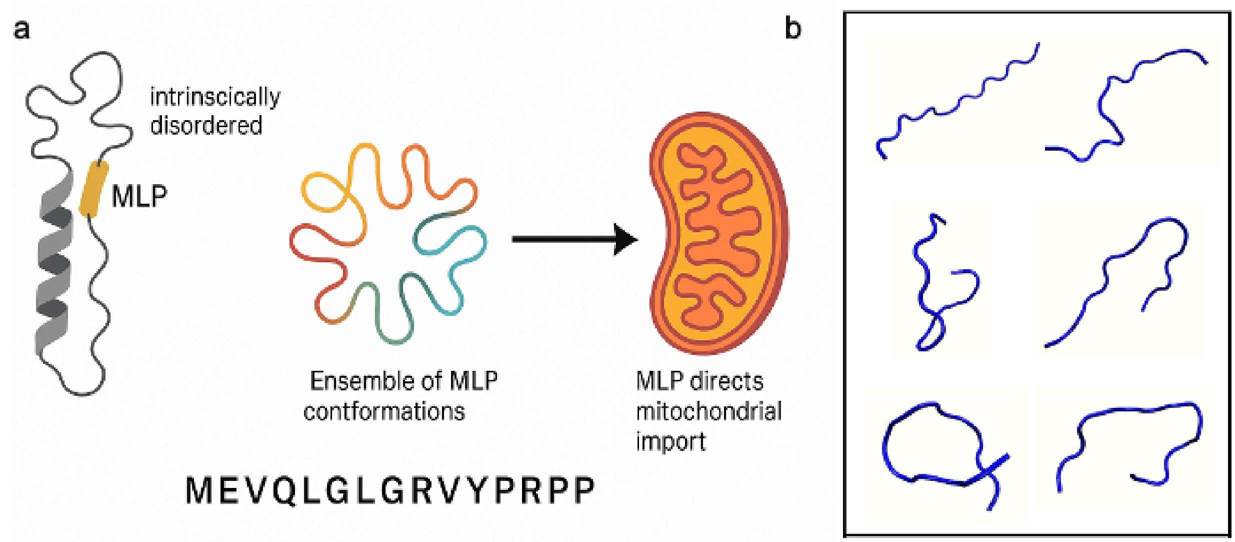


**Figure S1:** a) Schematic overview of the biological role of the mitochondrial localization peptide (MLP). b) Representative starting conformations generated for wild-type MLP following optimization of an initially linear structure. The conformations shown are illustrative and not exhaustive.

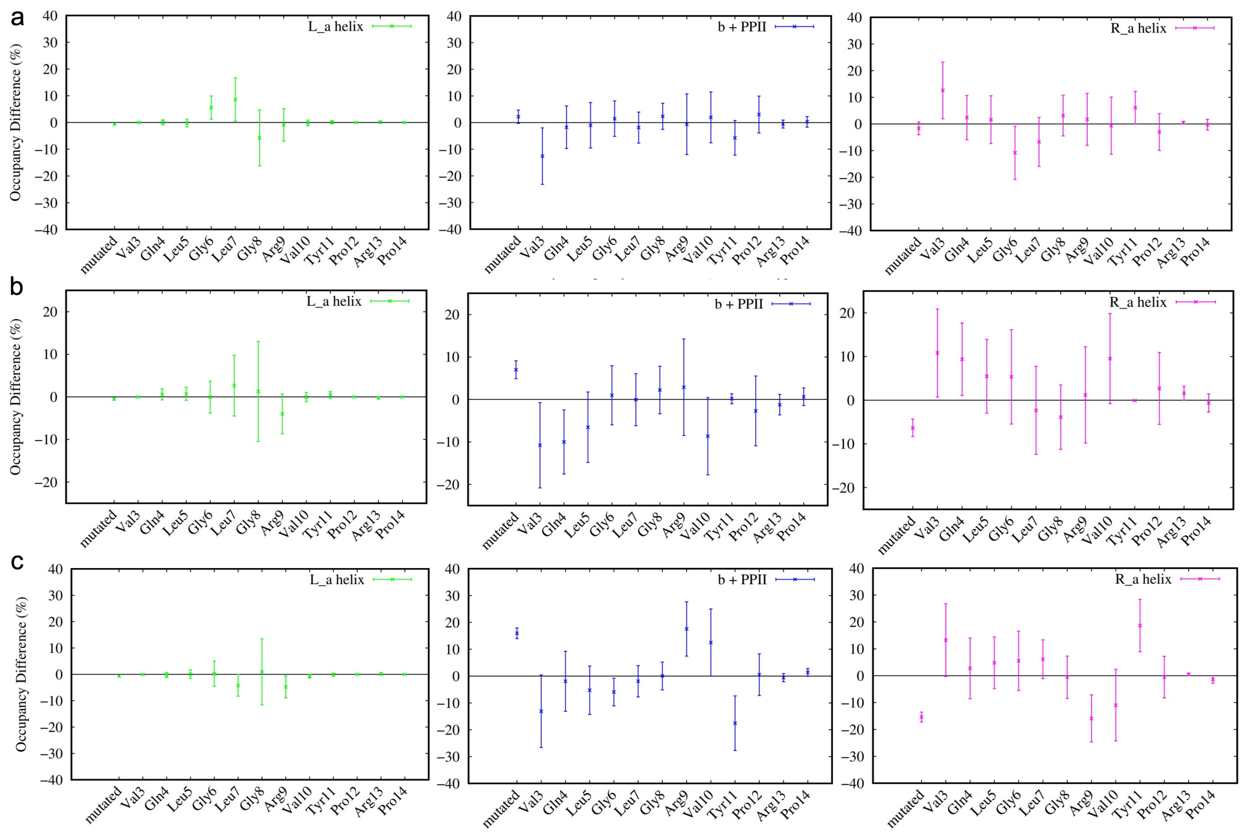


**Figure S2:** Plot showing secondary structural deviance of a) mlp e2c vs wild type, b) mlp e2d vs wild type and c) mlp e2f vs wild type. Occupancies are derived from Ramachandran plots as a ratio of how much time a given residue spends in each region and reflect an average of the 16 parallel simulations. Error bars reflect the appropriate spread of averages from both data sets.

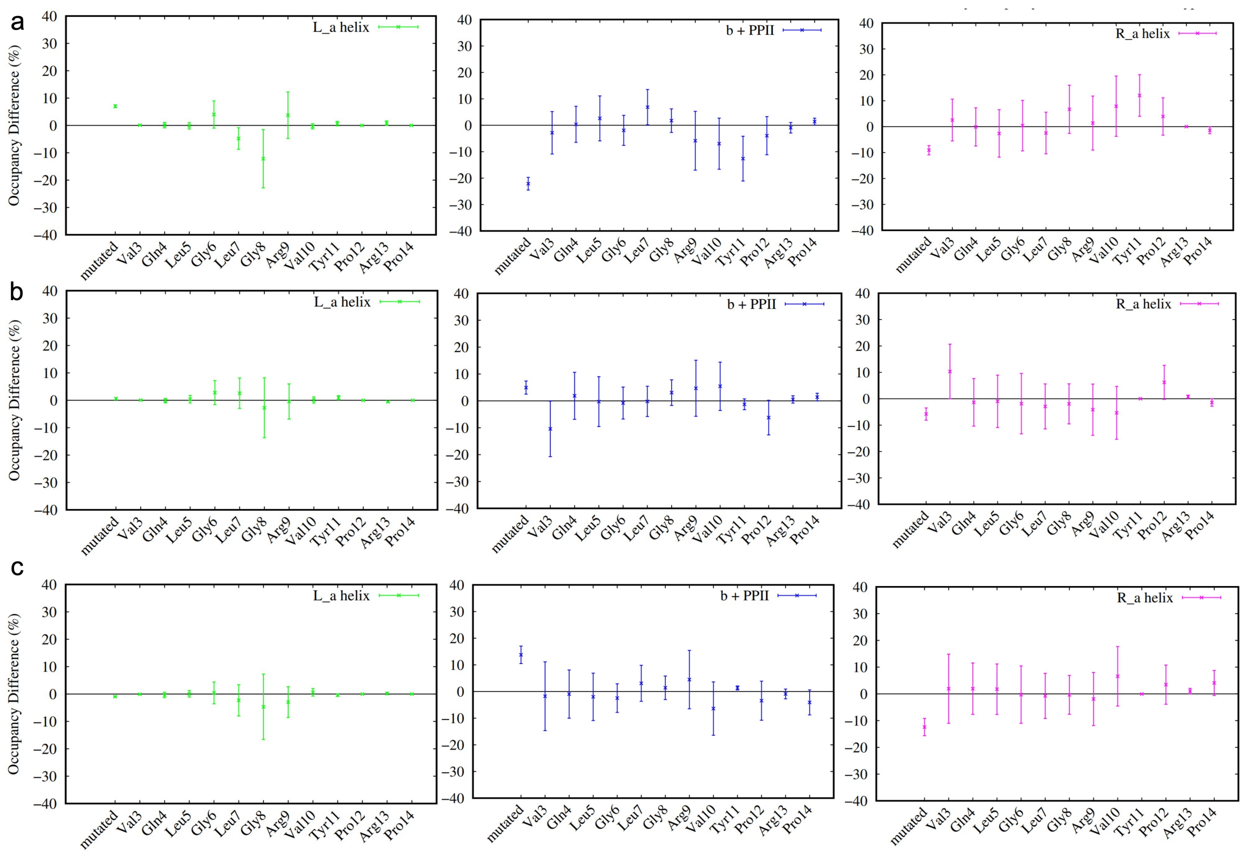


**Figure S3:** Plot showing secondary structural deviance of a) mlp e2g vs wild type, b) mlp e2h vs wild type and c) mlp e2i vs wild type. Occupancies are derived from Ramachandran plots as a ratio of how much time a given residue spends in each region and reflect an average of the 16 parallel simulations. Error bars reflect the appropriate spread of averages from both data sets.

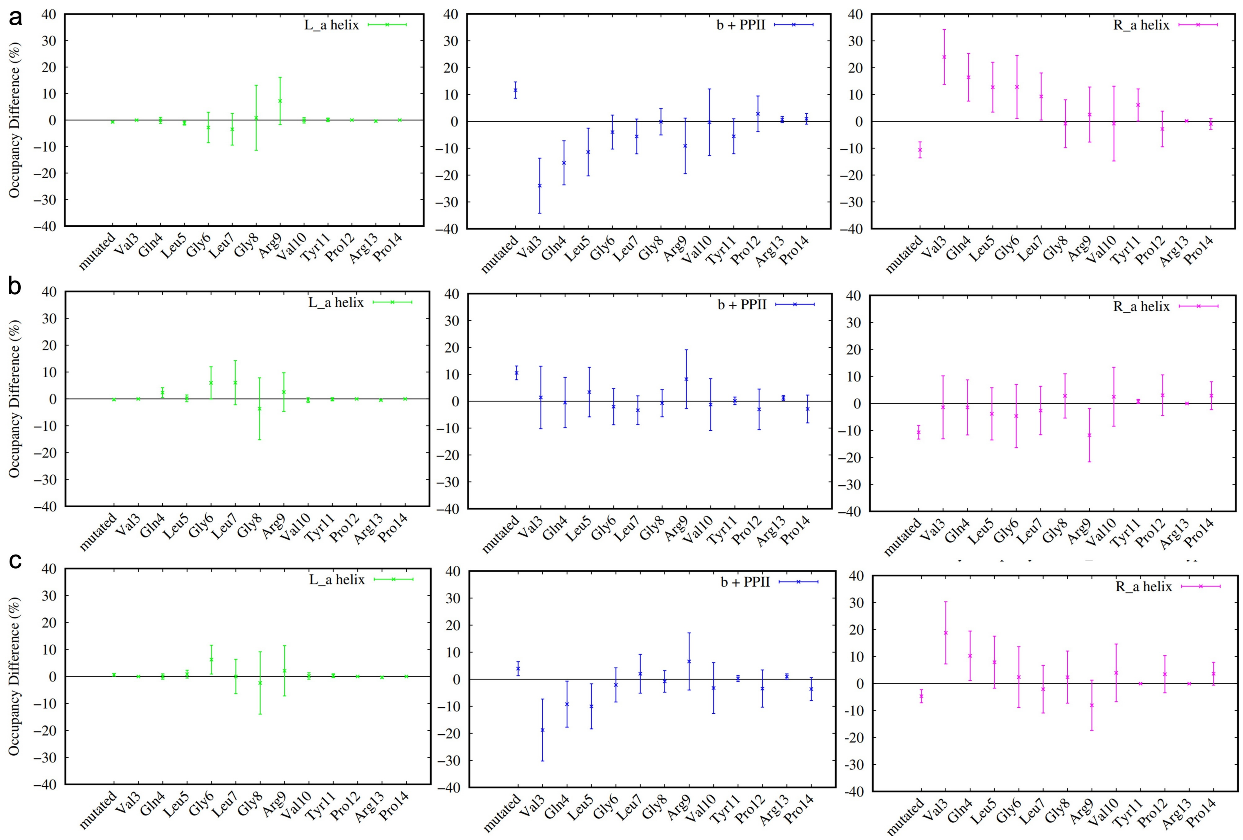


**Figure S4:** Plot showing secondary structural deviance of a) mlp e2l vs wild type, b) mlp e2m vs wild type and c) mlp e2n vs wild type. Occupancies are derived from Ramachandran plots as a ratio of how much time a given residue spends in each region and reflect an average of the 16 parallel simulations. Error bars reflect the appropriate spread of averages from both data sets.

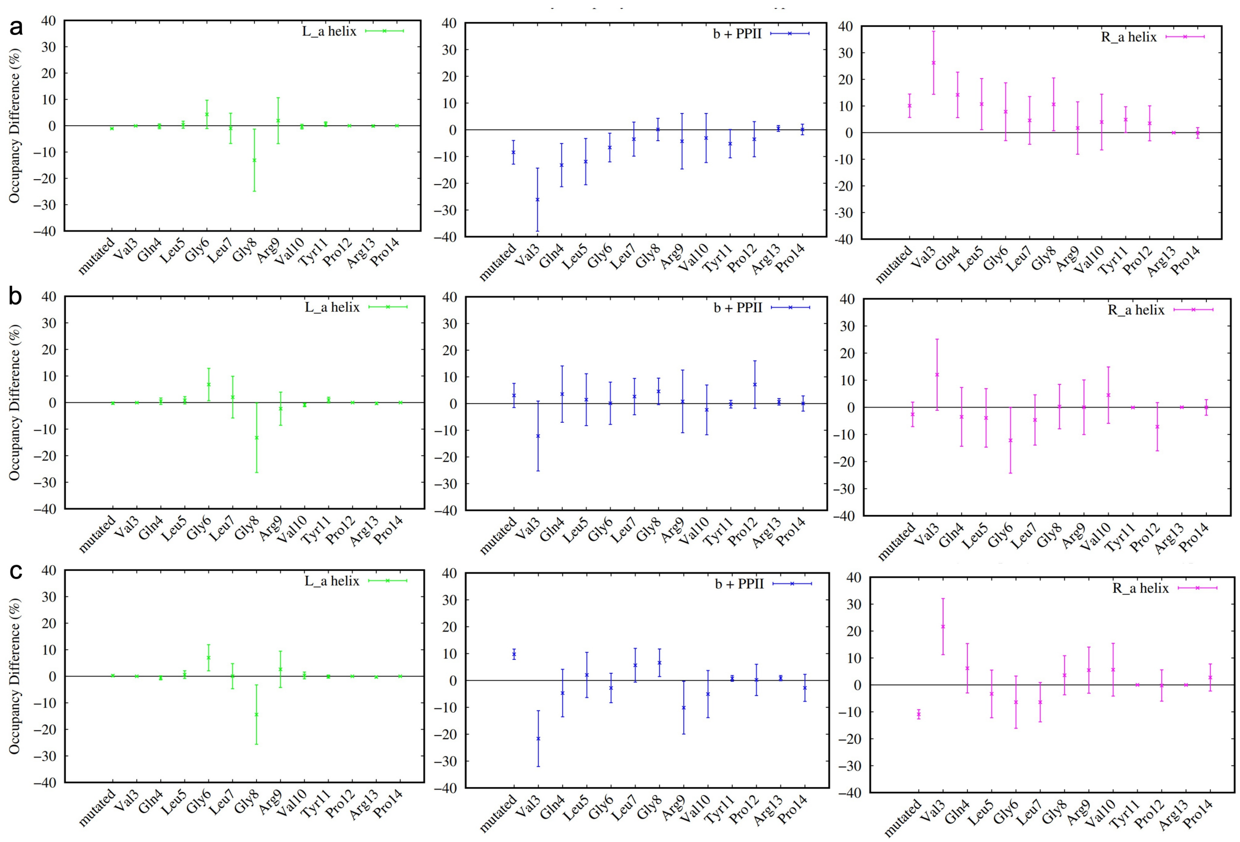


**Figure S5:** Plot showing secondary structural deviance of a) mlp e2p vs wild type, b) mlp e2r vs wild type and c) mlp e2s vs wild type. Occupancies are derived from Ramachandran plots as a ratio of how much time a given residue spends in each region and reflect an average of the 16 parallel simulations. Error bars reflect the appropriate spread of averages from both data sets.

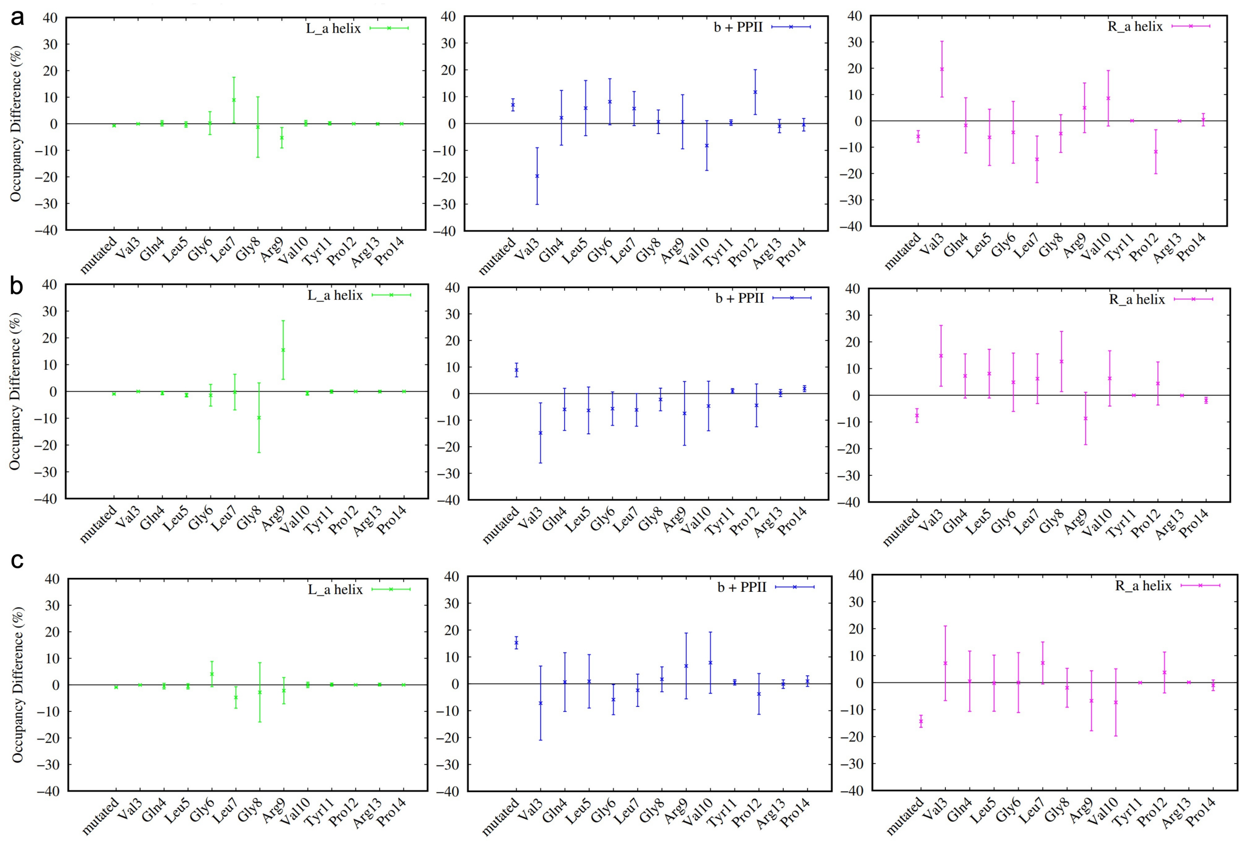


**Figure S6:** Plot showing secondary structural deviance of a) mlp e2t vs wild type, b) mlp e2v vs wild type and c) mlp e2w vs wild type. Occupancies are derived from Ramachandran plots as a ratio of how much time a given residue spends in each region and reflect an average of the 16 parallel simulations. Error bars reflect the appropriate spread of averages from both data sets.

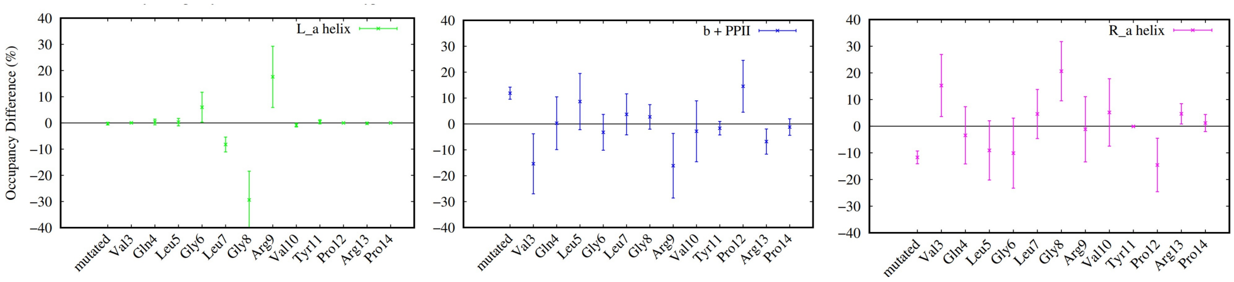


**Figure S7:** Plot showing secondary structural deviance of mlp e2y vs wild type.

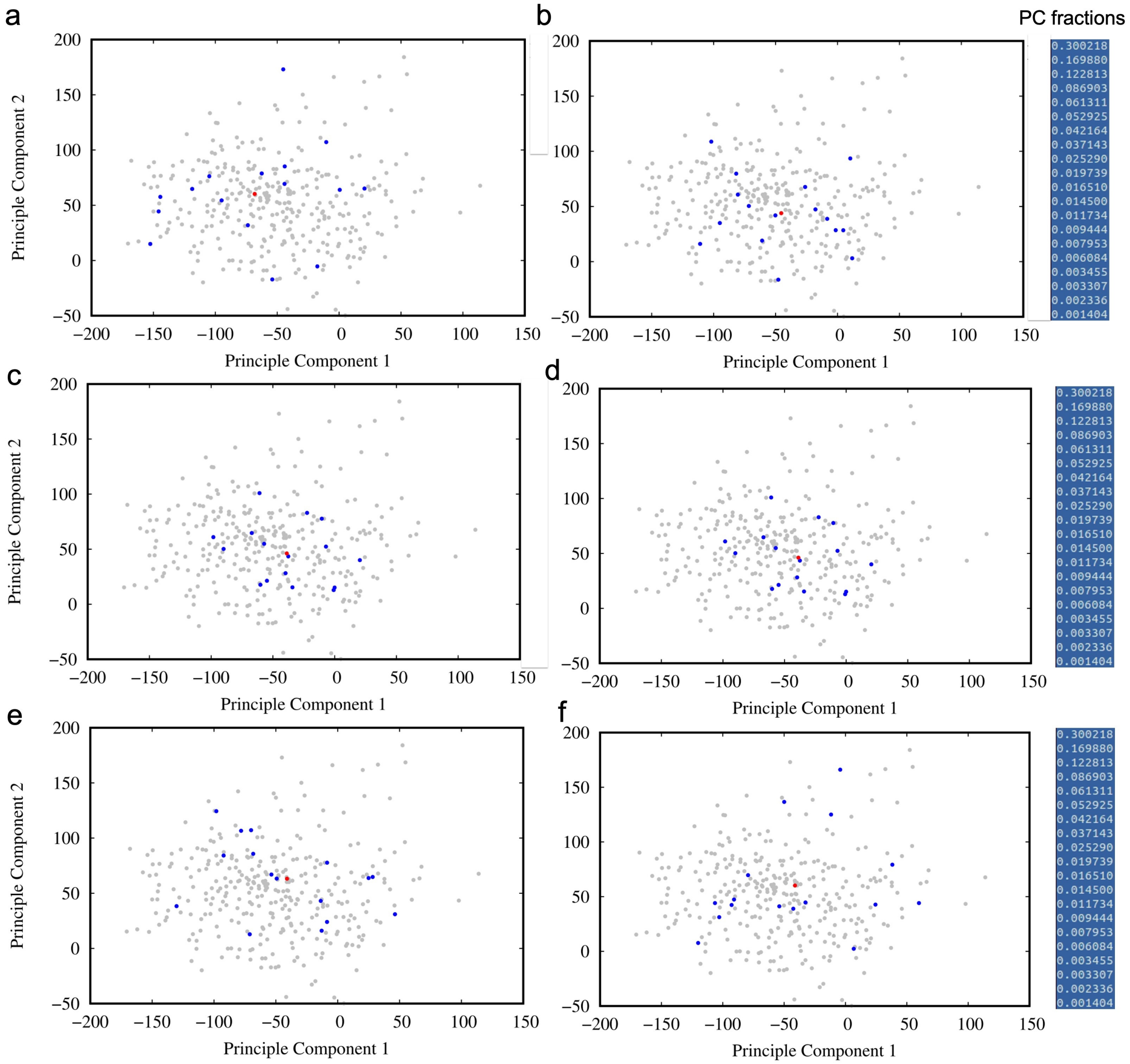


**Figure S8:** Principal component projection of secondary structural propensity data of a) mlp e2c, b) mlp e2d, c) mlp e2f, and d) mlp e2g e) mlk e2h f) mlk e2i. Each blue dot represents one of 16 parallel simulations, while red dot represents the averaged behavior over all 16 simulations. Grey dots show the spread of all remaining variants.

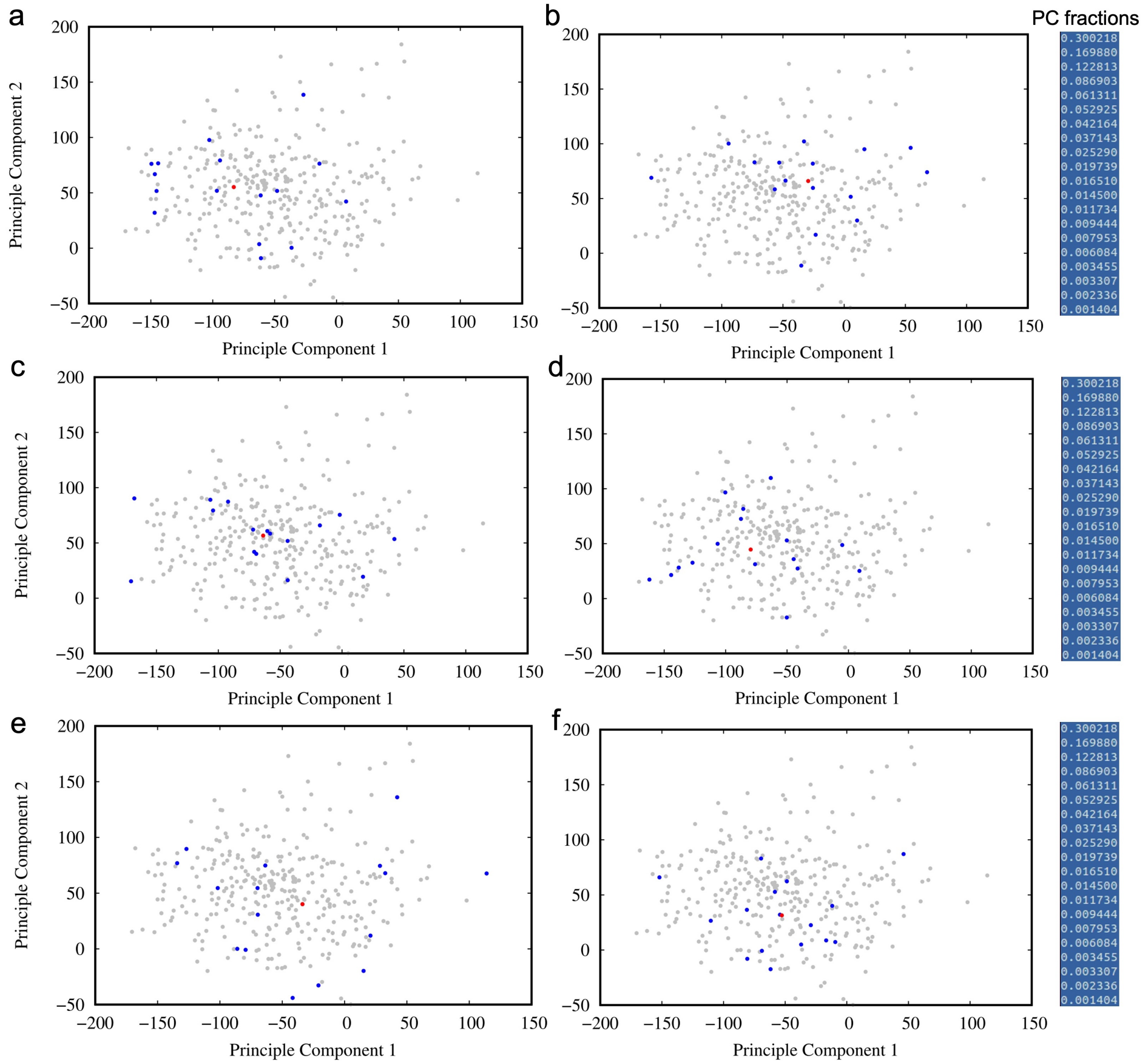


**Figure S9:** Principal component projection of secondary structural propensity data of a) mlp e2l, b) mlp e2m, c) mlp e2n, and d) mlp e2p e) mlk e2r f) mlk e2s. Each blue dot represents one of 16 parallel simulations, while red dot represents the averaged behavior over all 16 simulations. Grey dots show the spread of all remaining variants.

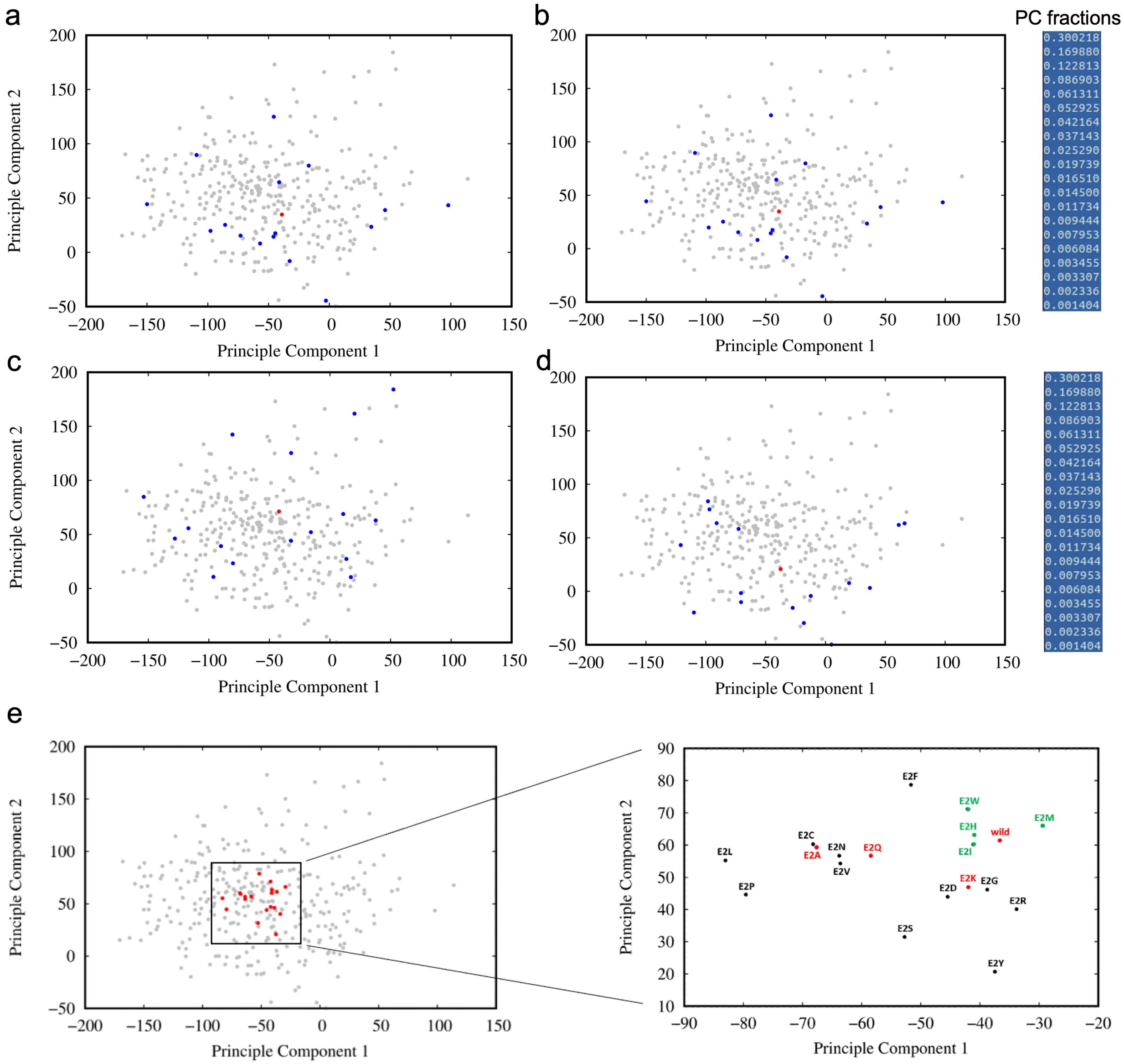


**Figure S10:** Principal component projection of secondary structural propensity data of a) mlp e2t, b) mlp e2v, c) mlp e2w, and d) mlp e2y e) mlk average. Each blue dot represents one of 16 parallel simulations, while red dot represents the averaged behavior over all 16 simulations. Grey dots show the spread of all remaining variants.

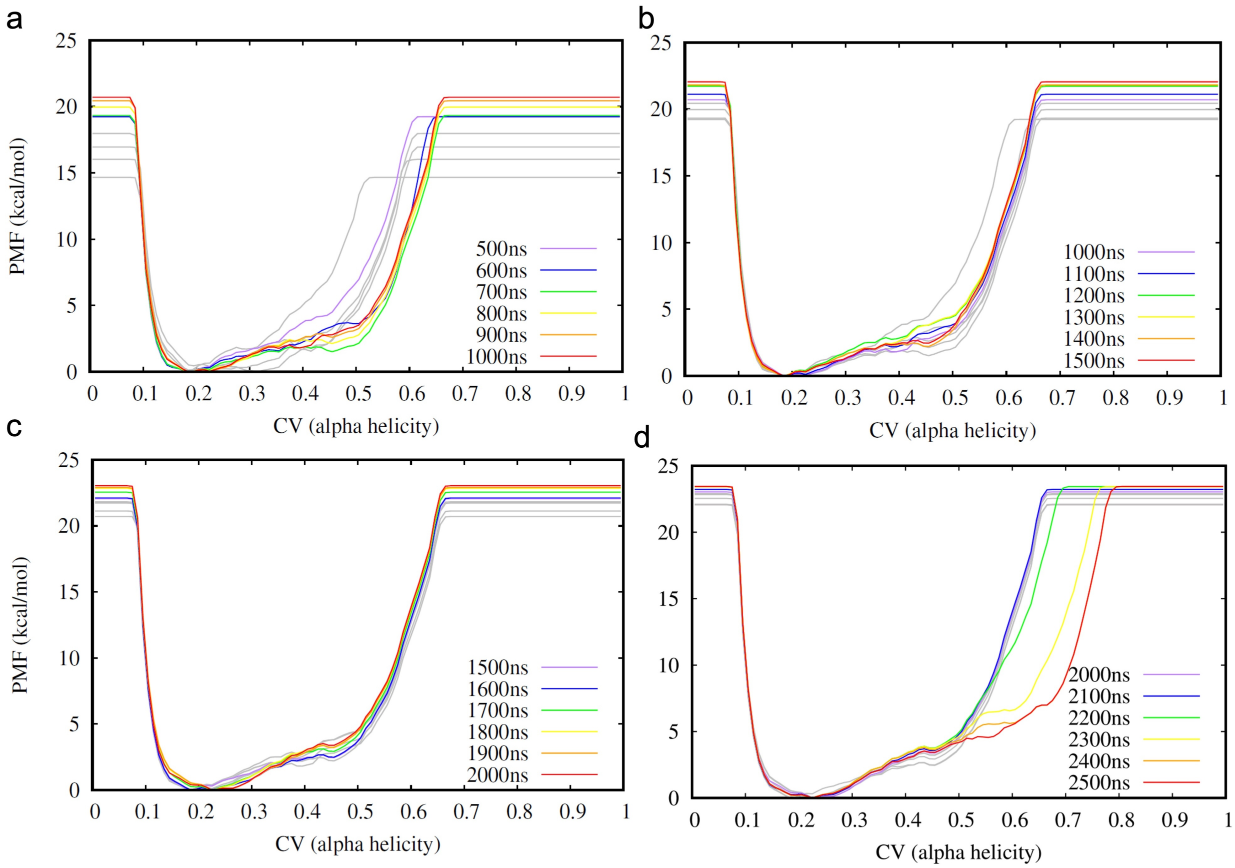


**Figure S11:** Time-evolving PMF for MLP e2a metadynamics for a) first 40%, b) first 60%, c) first 80%, and d) 100% of the 2.5 $\mu$s long MD simulation.

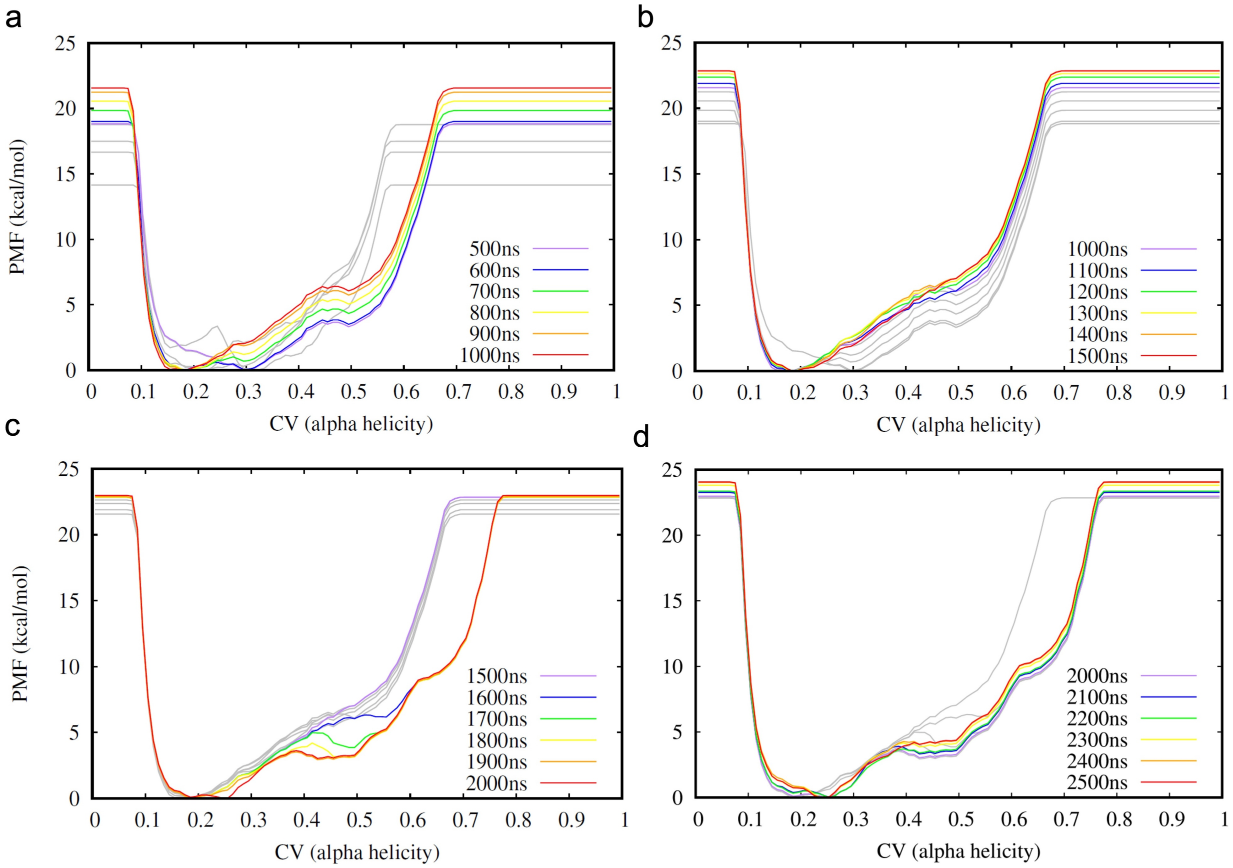


**Figure S12:** Time-evolving PMF for MLP e2k metadynamics for a) first 40%, b) first 60%, c) first 80%, and d) 100% of the 2.5 $\mu$s long MD simulation.

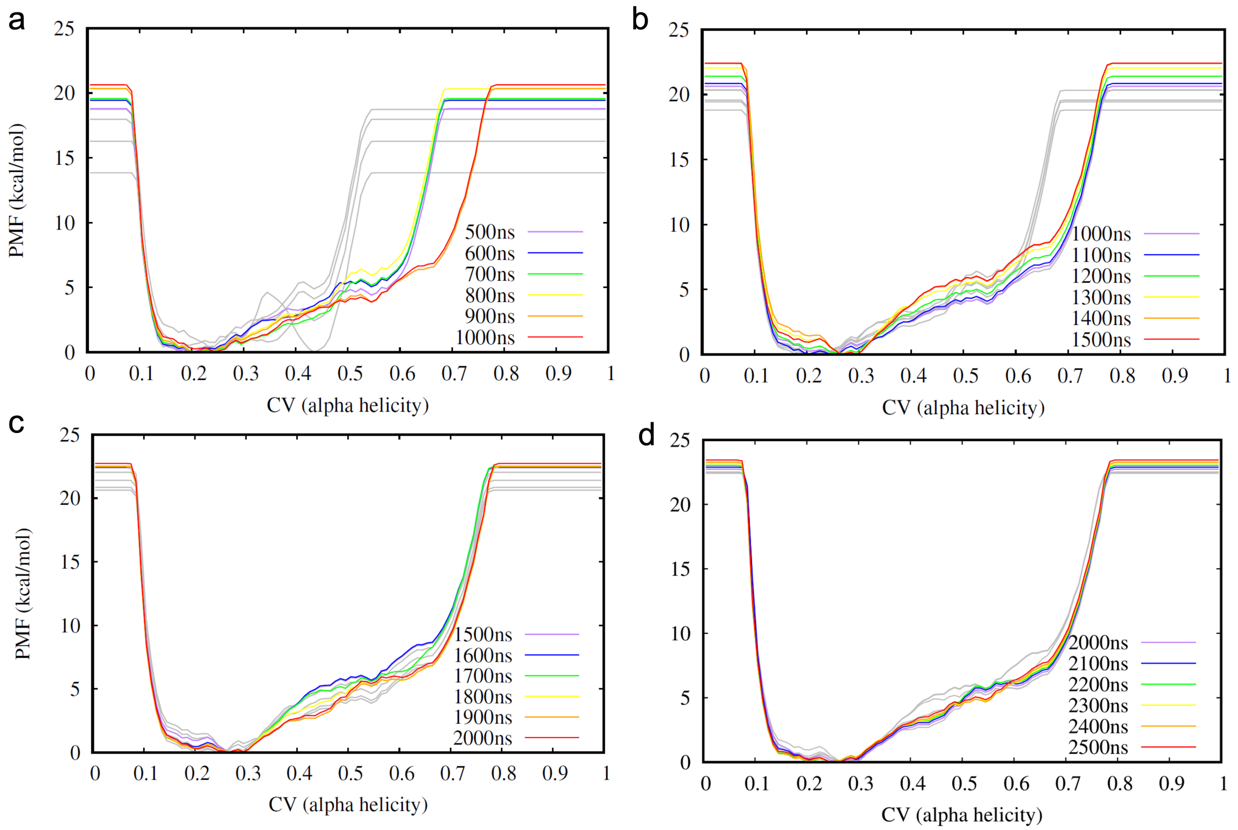


**Figure S13:** Time-evolving PMF for MLP e2q metadynamics for a) first 40%, b) first 60%, c) first 80%, and d) 100% of the 2.5 $\mu$s long MD simulation.

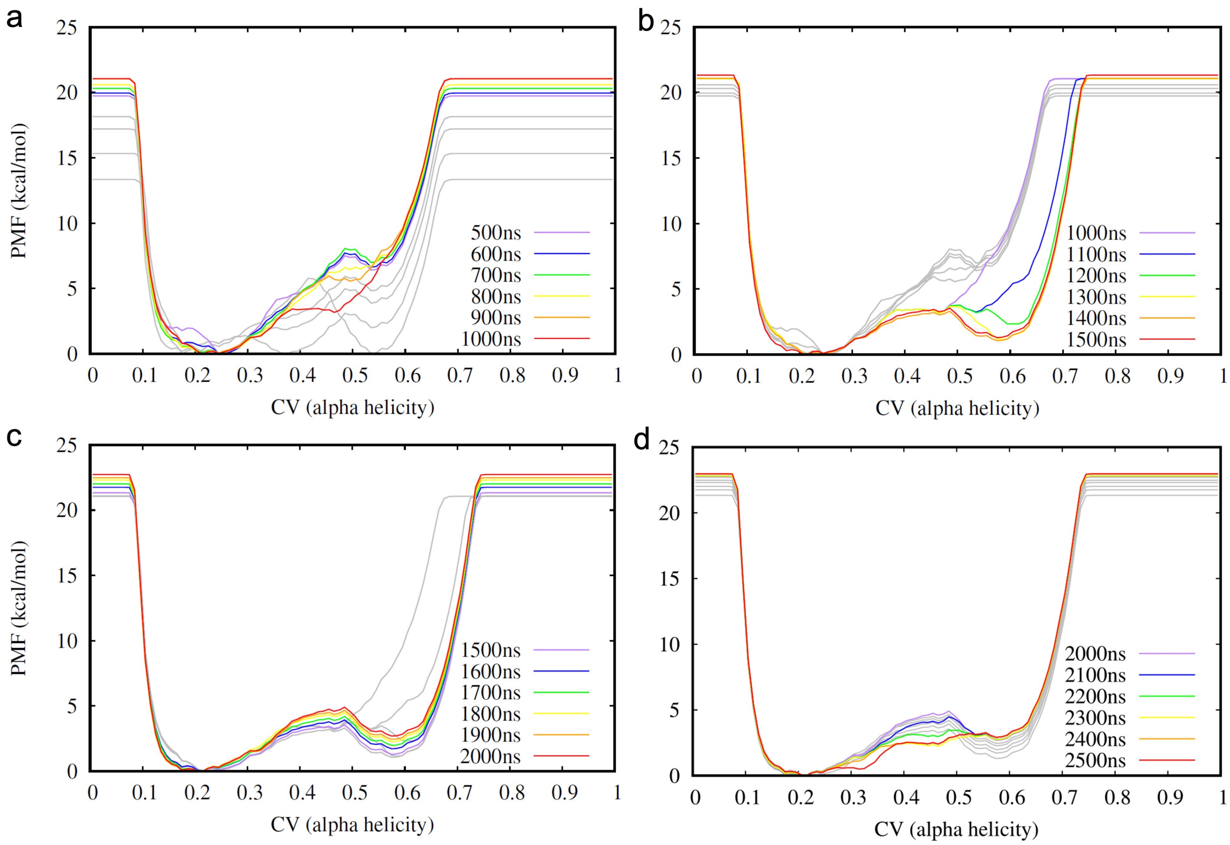


**Figure S14:** Time-evolving PMF for MLP wild-type metadynamics for a) first 40%, b) first 60%, c) first 80%, and d) 100% of the 2.5 $\mu$s long MD simulation.
